# Microtubules control cellular shape and coherence in amoeboid migrating cells

**DOI:** 10.1101/609420

**Authors:** Aglaja Kopf, Jörg Renkawitz, Robert Hauschild, Irute Girkontaite, Kerry Tedford, Jack Merrin, Oliver Thorn-Seshold, Dirk Trauner, Hans Häcker, Klaus-Dieter Fischer, Eva Kiermaier, Michael Sixt

## Abstract

Cells navigating through tissues face a fundamental challenge: while multiple cellular protrusions explore different paths through the complex geometry of an interstitial matrix the cell needs to avoid becoming too long or ramified, which might ultimately lead to a loss of physical coherence. How a cell surveys its own shape to inform the actomyosin system to retract entangled or stretched protrusions is not understood. Here, we demonstrate that spatially distinct microtubule (MT) dynamics regulate amoeboid cell migration by locally specifying the retraction of explorative protrusions. In migrating dendritic cells (DCs), the microtubule organizing center (MTOC) guides the path through a three dimensional (3D) interstitium and local MT depolymerization in protrusions remote from the MTOC triggers myosin II dependent contractility via the RhoA exchange factor Lfc. Depletion of Lfc leads to aberrant myosin localization, thereby causing two effects that rate-limit locomotion: i) impaired cell edge coordination during path-finding and ii) defective adhesion-resolution. Such compromised cell shape control is particularly hindering when cells navigate through geometrically complex microenvironments, where it leads to entanglement and ultimately fragmentation of the cell body. Our data demonstrate that MTs control cell shape and coherence by locally controlling protrusion-retraction dynamics of the actomyosin system.

---

How different cell types maintain their typical shape and how cells with a dynamic shape prevent loss of physical coherence is poorly understood. This issue becomes particularly critical in migrating cells, in which protrusion of the leading edge has to be balanced by retraction of the tail^1,2^ and where multiple protrusions of one cell often compete for dominance, as exemplified in the split pseudopod model of chemotactic migration^3^. The two prevalent models of how remote edges of mammalian cells communicate with each other are based on the sensing of endogenous mechanical parameters that in turn control the actomyosin system. In cell types that tightly adhere to substrates via focal adhesion complexes it has been proposed that actomyosin itself is the sensing structure and that adhesion sites communicate mechanically via actin stress fibers: when contractile stress fibers were pharmacologically, physically or genetically perturbed in mesenchymal cells, the cells lost their coherent shape and spread in an uncontrolled manner^4,5^. While communication via stress fibers is useful for adherent cells, it is unlikely to control the shape of amoeboid cells, which are often loosely or non-adherent and accordingly do not assemble stress fibers^6,7^. A second model suggests that lateral plasma membrane tension, which is thought to rapidly equilibrate across the cell surface, mediates communication between competing protrusions and serves as an input system to control actomyosin dynamics^8–11^. However, many amoeboid cells (such as DCs) are large and ramified^12^ and particularly when they are tightly embedded in 3D matrices it is questionable whether lateral membrane tension is able to equilibrate across the cell body^13^. Any alternative “internal shape sensor” would need to operate across the cellular scale and mediate communication between cell edges often more than hundred micrometers apart. Centrally nucleated MTs seem ideally suited for this task. We recently found that when leukocytes migrate through complex geometries, their nucleus acts as a mechanical gauge to lead them along the path of least resistance^14^. By spatial association with the nucleus, the MTOC and its nucleated MTs were involved in this navigational task, demonstrating that the positioning of the MTOC relative to the nucleus is critical for amoeboid navigation.

As cytoskeletal dynamics are notoriously difficult to visualize *in situ* or in physiological environments like collagen gels we used microfluidic pillar mazes^15^ as a reductionist setup that mimics some of the geometrical complexities of interstitial matrices while being accessible to imaging (Supplementary Fig. 1a-c). Within these devices cells are confined between two adjacent surfaces intersected by pillars of variable spacing and exposed to soluble gradients of the chemokine CCL19. CCL19 polarizes and directionally guides DCs and within the organism this leads them via lymphatic vessels towards the center of the draining lymph node^16^. To track MT plus ends we generated precursor cell lines stably expressing end binding protein 3 fused to mCherry (EB3-mCherry) and differentiated them into DCs. During time lapse imaging the MTOC appeared as the brightest spot radiating MT plus ends, indicating that MTs nucleate almost exclusively at the MTOC which is mainly located behind the nucleus (Fig. 1a, Supplementary Fig. 1d-f). When cells navigated through the pillar maze, the MTOC moved in a remarkably straight line up the chemokine gradient although transient lateral protrusions regularly explored alternative paths between the pillars (Fig. 1b). Notably, MT plus ends vanished from lateral protrusions that later became retracted (Supplementary Movie 1). To capture MT dynamics more quantitatively we imaged DCs migrating along chemokine gradients when confined under a pad of agarose^17^ where the flattened morphology allows faithful tracing of fluorescent signal (Supplementary Figure 1a, d). Here, cells migrate persistently and are stably segregated into a protruding leading edge and a retracting trailing edge (Fig. 1c). Visualization of the MT binding domain of ensconsin (EMTB) revealed long lived MTs at the leading lamellipodium while MT dynamics were increased at the trailing edge, exhibiting higher frequencies of shrinkage events compared to front directed filaments (Fig. 1d, Supplementary Fig. 1g and Supplementary Movie 2). These observations demonstrate that MT depolymerization is associated with cellular retraction.

**Figure 1.**
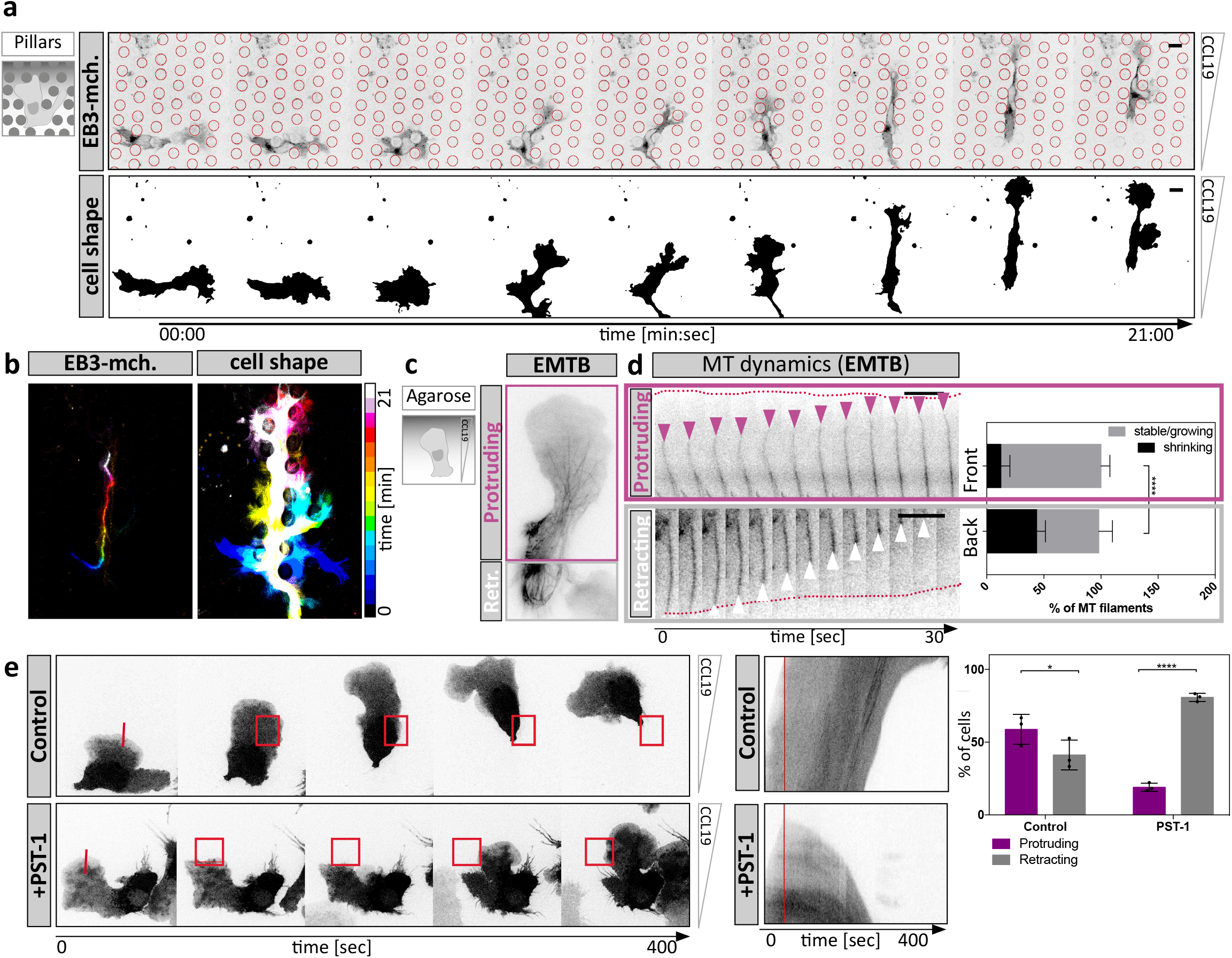
The microtubule organizing center acts as a pathfinder by coordinating protrusion dynamics. **a**, DC migrating within a pillar array. Upper panel shows EB3-mCherry expression profile. Lower panel outlines dynamic cell shape changes. Scale bar, 10μm. **b**, Time projection of image sequence shown in **a**. Left panel indicates MTOC position over time. Right panel outlines formation of multiple explorative protrusions over time. **c**, EMTB-mCherry expressing DC migrating under a pad of agarose. Purple box indicates protrusive cell front, whereas grey boxed area denotes contractile trailing edge. **d**, MT dynamics during directed migration. Growth (purple arrowheads) and shrinkage (white arrowheads) frequencies of individual MT filaments (according to EMTB labelling) were assessed in protrusive (front, purple box) vs. contractile (back, grey box) areas of the same migratory cell. Growth events and catastrophes ≥1μm were tracked for n = 10 filaments in the respective region of N = 8 cells. Mean ± S.D. **** p ≤ 0.0001. Scale bar, 5μm. **e**, Time-lapse sequence of control or PST-1 treated cells, which were locally photo-activated (red box) during migration under agarose (left panels). Middle panels display kymograph analysis of the photo-activated area shown on the left (red line). The time point of photo-activation is shown in red. Right panel: Frequency of local retractions upon photo-activation of control or PST-1 treated DCs during migration (n = 26 cells per condition ± S.D. from N = 3 experiments). * P ≤ 0.05, **** P ≤ 0.0001.

To test for a causal relationship between MT depolymerization and retraction we devised a photo-pharmacological approach to depolymerize MTs in migratory cells with spatiotemporal control. We used Photostatin-1 (PST-1), a reversibly photo-switchable analogue of combretastatin A-4, which can be functionally toggled between the active and inactive state by blue and green light, respectively^18^. To validate the approach, we locally activated the drug under simultaneous visualization of MT plus ends using EB3-mCherry. We found that local photo-activation triggered almost instantaneous disappearance of the EB3 signal in the presence but not in the absence of Photostatin (Supplementary Fig. 1h), indicating rapid stalling of MT polymerization. Local depolymerization in protruding areas of the cell led to consistent collapse of the illuminated protrusion and subsequent re-polarisation of the cell (Fig. 1e, Supplementary Movie 3). This response was only observed in the presence of Photostatin, while in the absence of the drug cells were refractory to illumination. These data demonstrate a causal relationship between MT depolymerization and cellular retraction. This effect can act locally within a cell, raising the possibility that MTs coordinate subcellular retractions when navigating through geometrically complex environments such as collagen gels or a physiological interstitium.

To directly address the impact of the MT cytoskeleton on the coordination of competing protrusions, we used a microfluidic setup in which DCs migrate in a straight channel towards a junction where the channel splits into four paths. In this setup DCs initially insert protrusions into all four channels before they retract all but one protrusion and thereby select one path along which they advance (Figure 2a, Supplementary Figure 1a). Global depletion of MTs using nocodazole (Supplementary Fig. 2a) led to uncoordinated protrusion dynamics and resulted in cell entanglement due to defective retraction of lateral protrusions (Figure 2b). Frequently, cells lost cytoplasmic coherence when competing protrusions continued to migrate up the chemokine gradient until the cell ruptured (Figure 2c, Supplementary Movie 4). Similarly, cells migrating *in situ* failed to reach lymphatic vessels (Supplementary Fig. 2b) and cells migrating in collagen gels lost cytoplasmic coherence and fragmented upon nocodazole treatment (Supplementary Fig. 2c, Supplementary Movie 5). In contrast to complex environments such as bifurcating channels and collagen gels, MT depolymerization did not affect cell coherence and general migratory capacity in linear microfluidic channels. In such geometrically simple environments where uniaxial polarity is externally enforced and where there is no competition of multiple protrusions, nocodazole merely caused cells to switch direction more frequently than untreated cells (Fig. 2d, e, Supplementary Movie 6, Supplementary Fig. 2d-g).

**Figure 2.**
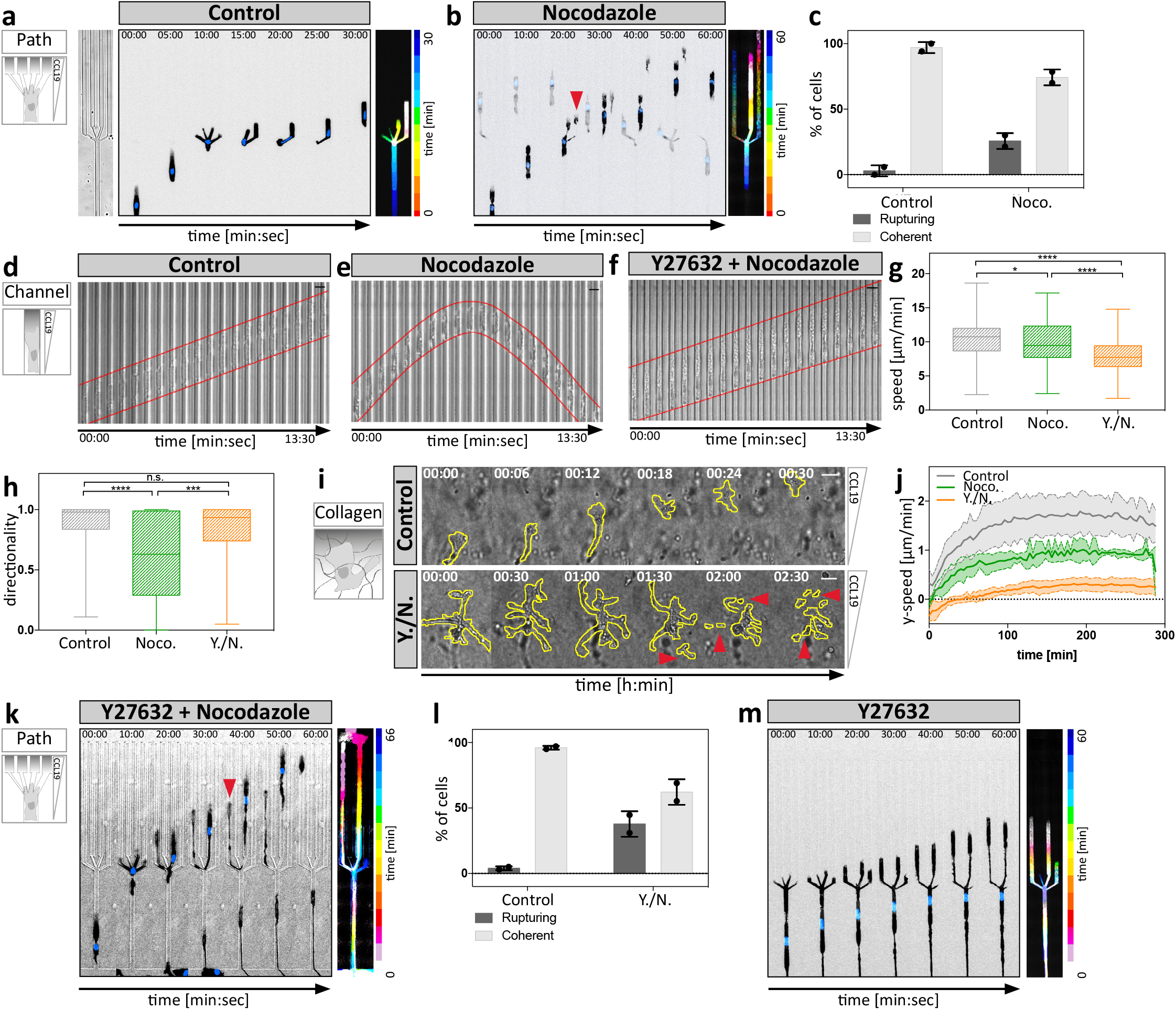
Microtubules coordinate protrusion dynamics via the contractile module. **a**, Left panel outlines channel geometry of path choice device. Middle panel: Lifeact-GFP expressing DC migrating within a path choice device of 5μm height. Time projection of signal distribution is shown in right panel. Scale bar, 10μm. **b**, Nocodazole-treated Lifeact-GFP expressing cell migrating within a path choice device. Note that the cell extends elongated protrusions into different channels. Red arrowhead denotes a cell rupturing event during the decision making process. Right panel outlines migratory behavior of cell and cellular fragment over time. **c**, Frequency of cell rupturing events during path choice decision. Control: n = 43 cells; nocodazole: n = 44 cells of N = 2 experiments. **d**, Time lapse sequence of a cell migrating within a linear microchannel of 5μm height. **e**, Nocodazole-treated cell migrating in the same configuration as in **d. f**, Cell treated with a combination of Y27632 plus nocodazole migrating as shown in **d**. Scale bar, 10μm. **g**, Migration speed of control, nocodazole- (Noco.) treated or double-treated (Y./N.) cells using Y27632 and nocodazole within microchannels (n = minimum of 74 cells per condition from N = 4 experiments). Boxes extend from 25th to 75th percentile. Whiskers span minimum to maximum values. **** P ≤ 0.0001. **h**, Directionalities of control, nocodazole- (Noco.) treated or double-treated (Y./N.) cells using Y27632 and nocodazole within microchannels (n = minimum of 74 cells per condition from N = 4 experiments). Boxes extend from 25th to 75th percentile. Whiskers span minimum to maximum values. **** P ≤ 0.0001, **** P ≤ 0.0001. **i**, DCs migrating within a collagen gel either non-treated (control) or double-treated with Y27632 and nocodazole (Y./N.). Note the different time intervals per condition. Red arrowheads indicate loss of cellular coherence in the double-treated cell. Scale bars, 10μm. **j**, Automated analysis of y-directed speed of non-treated, nocodazole-treated (Noco.) or double-treated cells using Y27632 and nocodazole (Y./N.). Plot shows mean migration velocities over time ± S.D. from N = 4 experiments. **k**, Lifeact-GFP expressing DC double-treated with Y27632 plus nocodazole (Y./N.) migrating as in **a**. Red arrowhead denotes a cell rupturing event during the decision making process. Right panel outlines migratory behavior of cell and cellular fragment over time. **l**, Frequency of cell rupturing events during path choice decision. Control: n = 40 cells; Y./N.: n = 80 cells of N = 2 experiments. **m**, Lifeact-GFP expressing DC treated with Y27632 migrating as in **a**. Note the extended protrusions reaching far into separate channels without generating a productive decision within the indicated time. Time projection of signal distribution is shown in right panel.

As demonstrated in other cell types, nocodazole treatment triggered a global increase of RhoA activity and myosin light chain (MLC) phosphorylation^19,20^ (Supplementary Fig. 2h, i) and pharmacological inhibition of the effector kinase Rho-associated protein kinase (ROCK) by Y27632 reverted this effect. Accordingly, in linear channels nocodazole-induced directional switching was reverted by additional ROCK inhibition (Fig. 2f-h, Supplementary Movie 6), indicating that an important function of MTs is to both control the activity and to stabilize the localization of the cell’s contractile module. As expected, ROCK inhibition failed to rescue cell integrity and locomotion after MT depletion in complex environments (Figure 2i-l, Supplementary Movie 5). Here, contractility is rate limiting for locomotion and ROCK inhibition alone caused the cells to entangle (Fig. 2m). Together our data add evidence that MTs act upstream of the contractile module and that actomyosin contractility is locally coordinated by MT depolymerisation, which effectively coordinates competing protrusions when cells migrate through complex environments.

One established molecular link between MT depolymerization and actomyosin contraction is the MT-regulated RhoA guanine nucleotide exchange factor (GEF) Lfc, the murine homologue of GEF-H1^21^. When Lfc is sequestered to MTs it is locked in its inactive state and only upon release from MTs it is targeted to membrane associated sites where it becomes active and triggers actomyosin contraction^22,23^. To test whether Lfc is involved in coordinating protrusions we knocked out *Arhgef2*, the Lfc encoding gene in mice (Supplementary Fig. 3a-f) and placed mature DCs into bifurcating microfluidic channels (Fig. 3a, b, Supplementary Fig. 4a). In line with the finding that Lfc mediates between MTs and myosin II (Supplementary Fig. 4b, c, d), Lfc^-/-^ DCs showed increased passage times due to defective retraction of supernumerary protrusions (Fig. 3c, d) although showing no differences in MT organization (Supplementary Fig. 4e-g). Similar to nocodazole-treated cells, Lfc^-/-^ DCs often advanced through more than one channel (Fig. 3b), ultimately resulting in auto-fragmentation into migratory cytoplasts (Fig. 3e, Supplementary Movie 7). When DCs migrated in straight channels and even when confronted with single constrictions, Lfc^-/-^ cells passed with the same speed and efficiency as wildtype cells (Fig. 3f, Supplementary Fig. 4h), demonstrating that neither locomotion nor passage through constrictions was perturbed but rather the coordination of competing protrusions. These data indicate that in complex 3D geometries, where the cell has to choose between different paths, MTs - via Lfc and myosin II - specify entangled protrusions for retraction.

**Figure 3.**
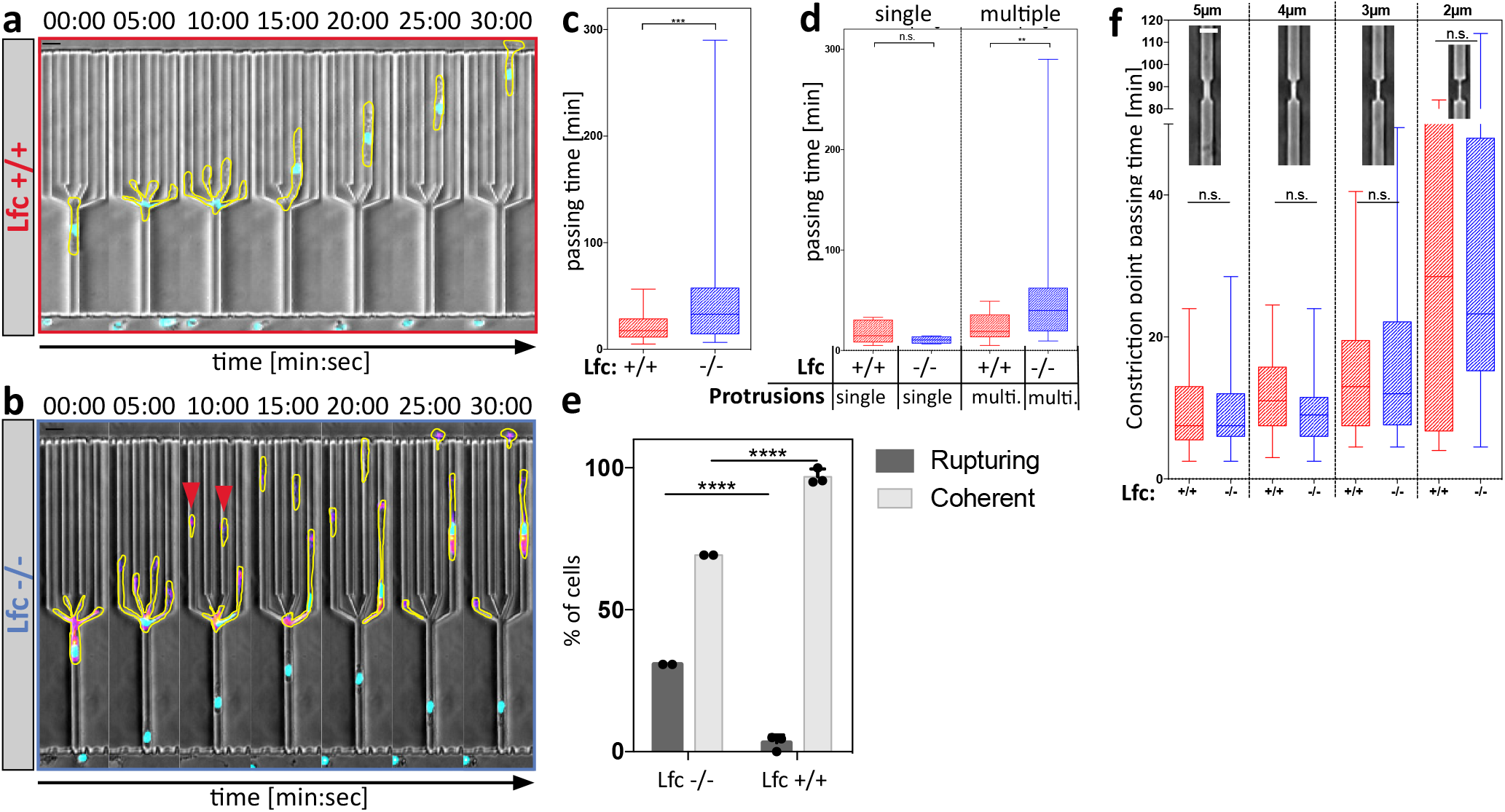
Microtubules mediate retraction of supernumerary protrusions via Lfc. **a**, Time lapse sequence of a wildtype littermate control cell migrating within a path choice device. Scale bar, 10μm. **b**, Time lapse sequence of a Lfc^-/-^ cell migrating within a path choice device. Red arrowheads denote multiple rupturing events of a single cell. Scale bar, 10μm. **c**, Junction point passing times of Lfc^+/+^ (n = 79 cells of N = 3 experiments) and Lfc^-/-^ (n = 49 cells of N = 2 experiments) DCs. Boxes extend from 25^th^ to 75^th^ percentile. Whiskers span minimum to maximum values. *** P ≤ 0.001. **d**, Junction point passing times depending on presence of single non-competing or multiple competing protrusions per cell of Lfc^+/+^ (n = 37 cells of N = 3 experiments) and Lfc^-/-^ (n = 46 cells of N = 2 experiments) DCs. Boxes extend from 25^th^ to 75^th^ percentile. Whiskers span minimum to maximum values. ** P ≤ 0.01. **e**, Frequency of cell rupturing events during path-choice decision of Lfc^+/+^ (n = 79 cells ± S.D. of N = 3 experiments) and Lfc^-/-^ (n = 52 cells ± S.D. of N = 2 experiments) DCs. **f**, Migration of DCs within straight, single constriction-containing microchannels. Graphs show constriction point passing times of Lfc^+/+^ (n = 114 cells of N = 3 experiments) and Lfc^-/-^ (n = 195 cells of N = 3 experiments) DCs. Boxes extend from 25^th^ to 75^th^ percentile. Whiskers span minimum to maximum values.

To determine how the Lfc pathway affects the contractile module we dynamically visualized MLC localization in Lfc^-/-^ and wildtype DCs migrating under agarose towards CCL19 gradients. While MLC was largely excluded from the leading lamellipodium, two distinct pools were detectable in the cell body of wildtype cells: one at the trailing edge and one in the cell center, at the base of the lamellipodium and around the nucleus (Fig. 4a, Supplementary Fig. 5a, b). Quantification of the distance between center of mass and center of MLC signal showed that migrating Lfc^-/-^ cells completely lost MLC polarization at the trailing edge, but maintained MLC in the cell center (Fig. 4b-d, Supplementary Fig. 5c, d and Supplementary Movie 8). The same distribution pattern was obtained by determining the localization of the active form of MLC (phospho-MLC) (Fig. 4e) and its effector protein moesin (Supplementary Fig. 5e, f) in fixed samples. This demonstrates that Lfc-deficiency leads to mislocalization of the contractile module and consequently to a loss of myosin II mediated contraction in peripheral protrusions. To address how this impacts overall locomotion, we next measured the migratory capacity of Lfc^-/-^ DCs under physiological conditions. *In situ* migration in explanted ear sheets showed that Lfc^-/-^ cells reached the lymphatic vessels later than control cells (Fig. 4f) and chemotaxis of Lfc^-/-^ DCs in collagen gels was substantially impaired (Fig. 4g, Supplementary Movie 9). When we measured cell lengths in 3D collagen gels, Lfc^-/-^ DCs were significantly elongated compared to control cells, indicating retraction defects (Fig. 4h).

**Figure 4.**
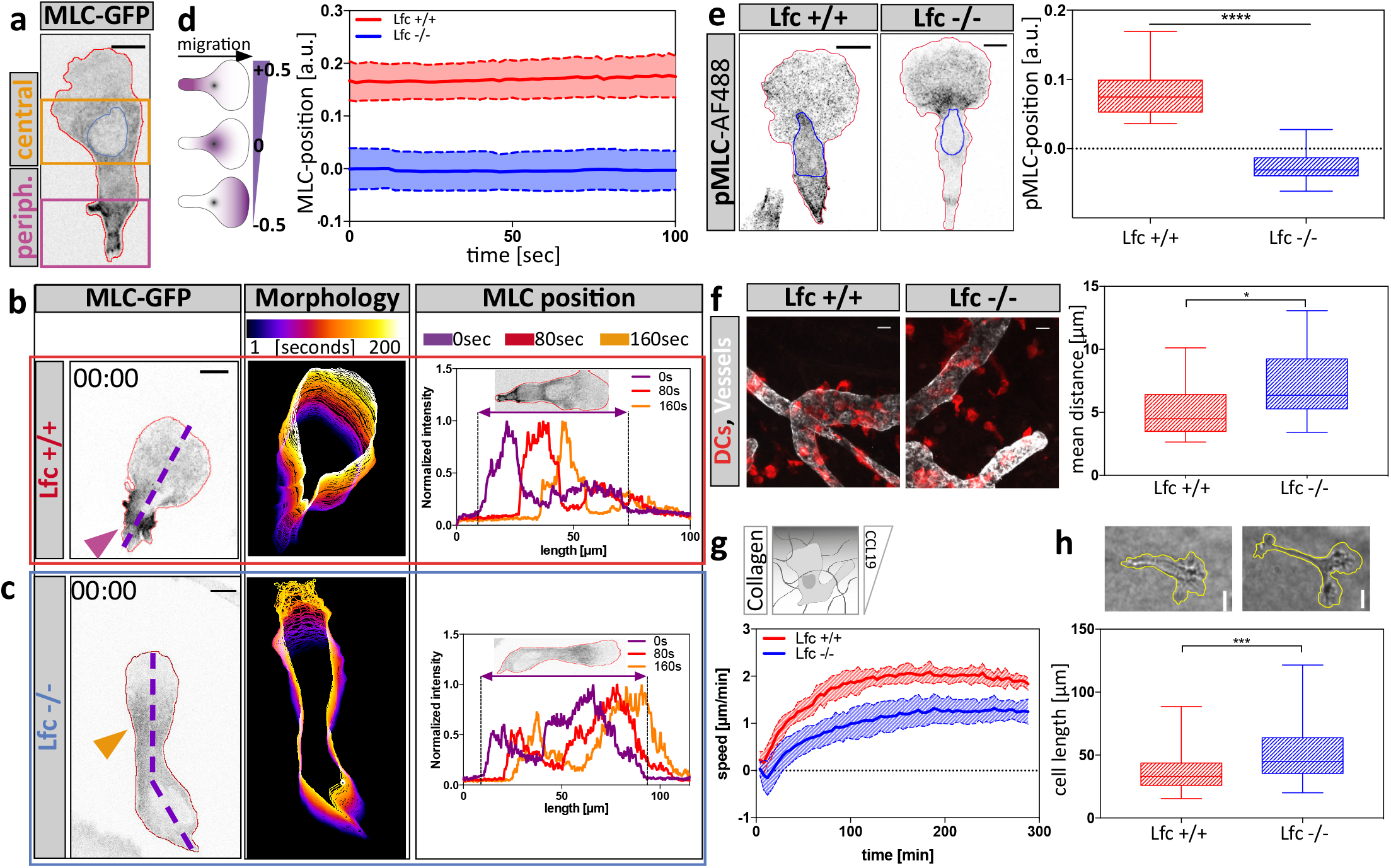
Lfc-dependent myosin accumulation controls cellular locomotion. **a**, A myosin light chain-GFP (MLC-GFP) expressing DC migrating under agarose along a soluble CCL19 gradient. Central- (orange box) and peripheral- (purple box) MLC accumulation is outlined. Scale bar, 10μm. MLC accumulation during migration under agarose in **(b)** wildtype- or **(c)** Lfc^-/-^ cells. Scale bar, 10μm. Middle panels indicate cell shapes over time. Right panels indicate mean MLC fluorescence distribution along the anterior-posterior polarization axis (dashed line) in 80sec intervals. **d**, Localization of MLC accumulation during directed migration of Lfc^+/+^ (red) and Lfc^-/-^ (blue) DCs. To account for differences in cell length the distance between cell center and MLC accumulation was normalized to cell length. Graph shows distance of n = 7 migratory cells per condition ± S.D. **e**, Localization of endogenous phospho-MLC(S19) in fixed migratory DCs (left panel). Right panel indicates position of MLC accumulation relative to cell length of n = 16 cells per condition from N = 4 experiments. Boxes extend from 25^th^ to 75^th^ percentile. Whiskers span minimum to maximum values. **** P ≤ 0.0001. Scale bar, 10μm. **f**, *In situ* migration of exogenous DCs on a mouse ear sheet. Lymphatic vessels were stained for Lyve-1, DCs with TAMRA respectively. Right panel indicates the mean distance of cells from lymphatic vessels. Per experiment two mouse ears with two fields of view were analyzed of N = 4 experiments. Boxes extend from 25^th^ to 75^th^ percentile. Whiskers span minimum to maximum values. * P ≤ 0.05. Scale bar, 100μm. **g**, Automated analysis of y-directed migration speed within a collagen network along soluble CCL19 gradient. Plot shows mean migration velocities over time ± S.D. from N = 7 experiments. **h**, Upper panels: Cell outlines of Lfc^+/+^ (left) and Lfc^-/-^ (right) DCs migrating within a collagen network along a soluble CCL19 gradient. Scale bar, 10μm. Lower panel shows lengths of cells migrating within a collagen network of n = 85 individual cells per condition from N = 4 experiments. Boxes extend from 25^th^ to 75^th^ percentile. Whiskers span minimum to maximum values. *** P ≤ 0.001.

In cells that employ an amoeboid mode of migration, defective retraction can not only stall locomotion by entanglement but might also lead to failed disassembly of integrin adhesion sites. We therefore tested the role of adhesion-resolution in under agarose assays, where, depending on the surface conditions, DCs can flexibly shift between adhesion-dependent and adhesion-independent locomotion^24^. Under adhesive conditions Lfc^-/-^ DCs were elongated compared to wildtype cells (Fig. 5a) and this elongation was lost when the migratory substrate at the bottom was passivated with polyethylene glycol (PEG) (Fig. 5b, e). When cells on adhesive surfaces were treated with nocodazole, wildtype cells shortened, as expected due to hypercontractility (Fig. 5c). Notably, Lfc^-/-^ DCs elongated even more upon treatment with nocodazole (Fig. 5c, lower panel), indicating that elimination of Lfc-mediated hypercontractility unmasked additional modes of MT-mediated length control. Elongation of Lfc^-/-^ cells by nocodazole was also largely absent on PEG-coated surfaces (Fig. 5d, f, Supplementary Movie 10). Importantly, not only morphological, but also migratory parameters were restored on passivated surfaces (Fig. 5g, h). Together, these data demonstrate that whenever DCs migrate in an adhesion-mediated manner, MTs control deadhesion and this is partially mediated via Lfc and myosin II. We conclude that MT depolymerization in peripheral regions of migrating DCs locally triggers actomyosin-mediated retraction via the RhoA GEF Lfc. Thereby MTs coordinate protrusion-retraction dynamics and prevent that the cell gets too long or ramified.

**Figure 5.**
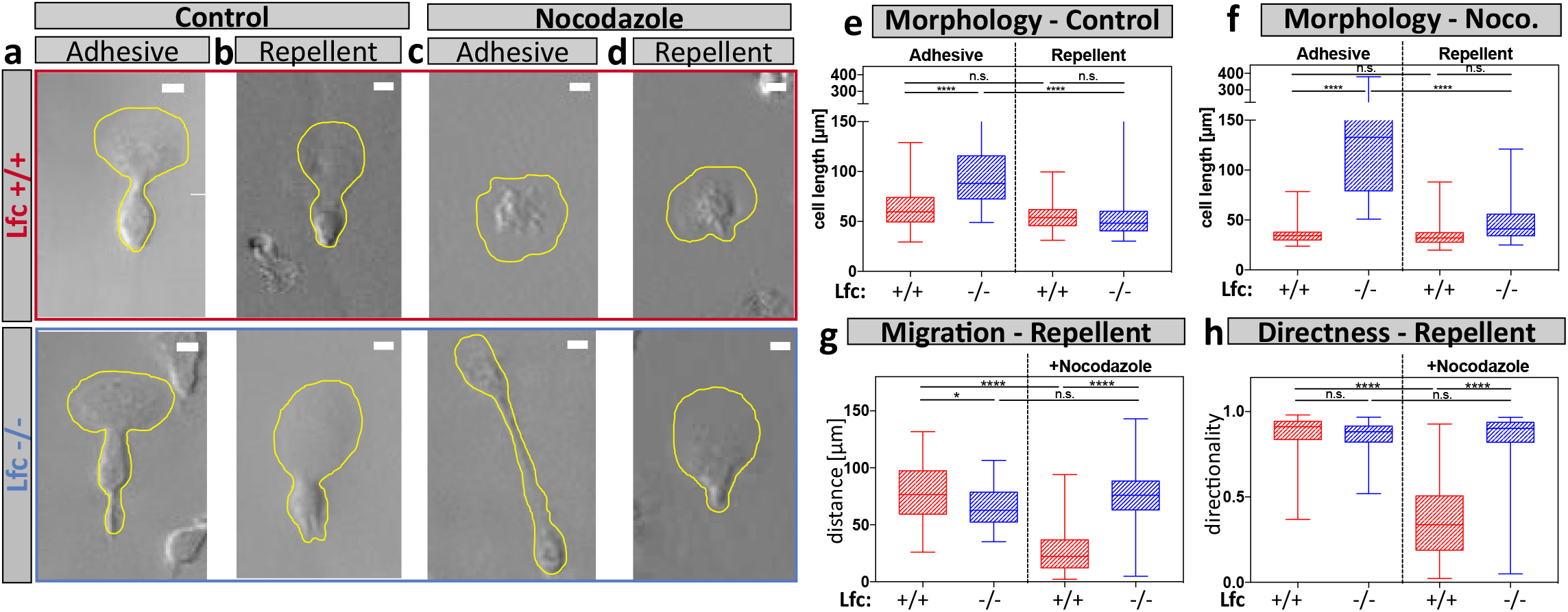
Lfc regulates microtubule-mediated adhesion resolution. Cell shape outlines of non-treated control cells migrating under agarose under adhesive **(a)** or repellent (PEG coated) **(b)** conditions. Cell shape outlines of nocodazole-treated cells migrating under agarose under adhesive **(c)** or repellent (PEG coated) **(d)** conditions. Upper panels show littermate control wildtype cells, lower panels show Lfc^-/-^ cells. Scale bars, 10μm. **e**, Cell lengths of non-treated control cells migrating under adhesive and repellent conditions (n = minimum of 80 cells per condition from N = 5 experiments). Boxes extend from 25^th^ to 75^th^ percentile. Whiskers span minimum to maximum values. **** P ≤ 0.0001. **f**, Cell lengths of nocodazole-treated cells migrating under adhesive and repellent conditions (n = minimum of 80 cells per condition from N = 5 experiments). Boxes extend from 25^th^ to 75^th^ percentile. Whiskers span minimum to maximum values. **** P ≤ 0.0001. **g**, Migration distance of Lfc^*+/+*^ and Lfc^-/-^ DCs migrating under agarose under non-adhesive (PEG coated) conditions of n = minimum of 80 cells per condition from N = 5 experiments. Cells were either non-treated or treated with nocodazole. Boxes extend from 25^th^ to 75^th^ percentile. Whiskers span minimum to maximum values. * P ≤ 0.05, **** P ≤ 0.0001. **h**, Directionalities of Lfc^+/+^ and Lfc^-/-^ DCs migrating under agarose under non-adhesive conditions (PEG). Cells were either non-treated or nocodazole-treated (n = minimum of 80 cells per condition from N = 5 experiments). Boxes extend from 25^th^ to 75^th^ percentile. Whiskers span minimum to maximum values. **** P ≤ 0.0001.

Although it is likely that multiple feedback loops signal between actin and MTs, we show that there is a strong causal link between local MT catastrophes and cellular retraction, with MTs acting upstream. This raises the key question how MT stability is locally regulated in DCs. Among many possible inputs (adhesion, chemotactic signals etc.) one simple option might be related to the fact that in leukocytes the MTOC is the only site where substantial nucleation of MTs occurs. In complex environments (like the pillar maze we devised) the MTOC of a DC moves a remarkably straight path, while lateral protrusions constantly explore the environment (Figure 1b). Hence, passage of the MTOC beyond an obstacle is the decisive event determining the future trajectory of the cell. Upon passage of the MTOC, sheer geometry might determine that all but the leading protrusion are cut off MT supply because MTs are too inflexible to find their way into curved, narrow and ramified spaces. Together, we propose that MTs serve as an internal explorative system of the cell that informs actomyosin whenever a peripheral protrusion locates too distant from the centroid and thereby initiates its retraction.

## Acknowledgments

The authors thank the Scientific Service Units of IST Austria for their excellent services. This work was supported by the European Research Council (ERC StG 281556 and CoG 724373), a grant from the Austrian Science Foundation (FWF) and the FWF DK ‘Nanocell’ to M.S. J.R was supported by ISTFELLOW funding from the People Programme (Marie Curie Actions) of the European Union’s Seventh Framework P (FP7/2007-2013) under REA grant agreement n° [291734] and an EMBO long-term fellowship (ALTF 1396-2014) co-funded by the European Commission (LTFCOFUND2013, GA-2013-609409). H.H. was supported by the American Lebanese Syrian Associated Charities (ALSAC).

## Materials and Methods

### Mice

All mice used in this study were bred on a C57BL/6J background and maintained at the institutional animal facility in accordance with the IST Austria ethics commission in accordance with the Austrian law for animal experimentation. Permission of all experimental procedures was granted and approved by the Austrian federal ministry of science, research and economy (identification code: BMWF-66.018/0005-II/3b/2012).

### Generation of Lfc^-/-^ mice

A cosmid containing the full genomic sequence of the gene that encodes Lfc (*Arhgef2*) was isolated from a 129 mouse genomic library with Lfc cDNA probes (106-630, 631-1057 and 1060-1478 bp) amplified by RT-PCR. The genomic DNA region between base pairs 1193-1477, coding for amino acids 351-445 in the DH domain and DH/PH domain interface was exchanged for a neomycin cassette flanked by LoxP sites. The targeting construct was linearized with *Not*I and electroporated into R1 ES cells. Homologous recombinants were selected in the presence of G418 (150 μg/ml) and gancyclovir (2 μM) and analyzed by Southern blotting. Positive embryonic stem cell clones were aggregated with eight cell-stage mouse embryos to generate chimeras. The resulting mice were genotyped by Southern blot and PCR. Primers (5′–CGGGGATCCATTCGGTTGTAA–3′) and (5′–AAGCGGCATGGAGTTCAGGA–3′) amplified a 365-bp fragment specific for the wild type allele, whereas primers (5′–AGAGTTCTGCAGCCGCCACACCA–3′) and 5′-GGTGGGGGTGGGGTGGGATTAGATA –3′) amplified a 500-bp fragment specific for the targeted allele. We refer to these mice as Lfc^-/-^ mice throughout the entire manuscript. Western blot analysis using a Lfc-specific antibody was performed to confirm that Lfc^-/-^ mice had no expression of Lfc protein. Mice were backcrossed to C57BL/6 background for more than 12 generations. Dendritic cells were generated from bone marrow isolated from littermates or age-matched wildtype and Lfc^-/-^ 8-12 week-old mice. Mice were bred and housed in accordance with institutional guidelines.

### Generation of immortalized hematopoietic progenitor reporter cell lines

Hematopoietic progenitor cell lines were generated by retroviral delivery of an estrogen-regulated form of HoxB8 as described recently ^25,26^. Briefly, bone marrow of 6-12 week of Lfc^+/+^ and Lfc^-/-^ mice was isolated and retrovirally transduced with an estrogen-regulated form of the HoxB8 transcription factor. After expansion of immortalized cells, lentiviral spin infection (1500g, 1h) was carried out in the presence of 8μg/ml Polybrene and the lentivirus coding for fluorescent expression construct of interest. Following transduction, cells were selected for stable virus insertion using 10μg/ml Blasticidin for at least one week. Cells expressing fluorescent reporter constructs were sorted using fluorescence-activated cell sorting (FACS Aria III, BD Biosciences) prior to migration experiments.

### Dendritic cell culture

Culture was started either from freshly isolated bone marrow of 6-12 week old mice with C57BL/6J background (wildtype, Lfc^-/-^, or Lifeact-GFP^27^ as described earlier^28^ or from stable hematopoietic progenitor cell lines after washing out estrogen. DC differentiation was induced by plating 2×10^6^ cells (bone marrow) or 2×10^5^ cells (progenitor cells) in complete media (RPMI 1640 supplemented with 10% Fetal Calf Serum, 2mM L-Glutamine, 100U/ml Penicillin, 100μg/ml Streptomycin, 50μM β-Mercaptoethanol) (all purchased from Invitrogen) containing 10% Granulocyte-Monocyte colony stimulating factor (GM-CSF, supernatant from hybridoma culture). To induce maturation, cells were stimulated overnight with 200ng/ml Lipopolysaccharide from *E.coli* 0127:B8 (Sigma) and used for experiments on days 9-10.

### *In situ* migration assay

Six to eight weeks old female C57BL/6J mice were sacrificed and individual ear sheets separated into dorsal and ventral halves as described previously^29^. Cartilage free ventral halves were incubated for 48h at 37°C, 5% CO_2_ with ventral side facing down in a well plate filled with complete medium. The medium was changed once 24h post-incubation-start. If indicated, pharmacological inhibitors were added to the medium. Ear sheets were fixed with 1% PFA followed by immersion in 0.2% Triton X-100 in PBS for 15min and three washing steps á 10min with PBS. Unspecific binding was prevented by 60min incubation in 1%BSA in PBS at room temperature. Incubation with primary rat-polyclonal antibody against LYVE-1 in combination with rat-polyclonal biotinylated anti-MHC-II antibody (both R&D Systems) was done for 2h at room temperature. After three times 10min washing with 1% BSA in PBS consecutive incubation using Alexa Fluor 488-AffiniPure F(ab’)_2_ fragment donkey anti-rat IgG (H+L) secondary antibody and streptavidin-Cy3 secondary antibody (both Jackson ImmunoResearch) was done. Samples were incubated 45min in first secondary antibody in the dark followed by 10min washing in 1% BSA in PBS and subsequent incubation with second secondary antibody. Samples mounted with ventral side up on a microscope slide, protected with a coverslip and stored at 4°C in the dark.

In order to determine the distance between the lymphatic vessels and DCs a mask was created by manually outlining lymphatic vessels depending on Lyve-1 staining and segmenting cells according to their fluorescence intensity. The distance between cells and lymphatic vessels was quantified using a custom-made Matlab script, which determines the closest distance from the segmented cells to the border of the lymphatic vessel binary image. Image borders were excluded from analysis.

### *In vitro* collagen gel migration assay

Custom made migration chambers were assembled by using a plastic dish containing a 17mm hole in the middle, which was covered by coverslips on each side of the hole. Three-dimensional scaffolds consisting of 1.73mg/ml bovine Collagen I were reconstituted *in vitro* by mixing 3×10^5^ cells in suspension with Collagen I suspension buffered to physiological pH with Minimum Essential Medium and Sodium Bicarbonate in a 1:2 ratio. To allow polymerization of Collagen fibers, gels were incubated 1h at 37°C, 5% CO_2_. Directional cell migration was induced by overlaying the polymerized gels with 0.63μg/ml CCL19 diluted in complete media (R&D Systems). To prevent drying-out of the gels, migration chambers were sealed with Paraplast X-tra (Sigma-Aldrich). The acquisition was performed in 60sec intervals for five hours at 37°C, 5% CO_2_. Detailed description of experimental procedure can be found elsewhere^30^.

### Analysis of y-displacement

Quantification of y-directed migration analysis of cell population was performed as described earlier ^31^. Briefly, raw data image sequences were background corrected and particles smaller and bigger than an average cell were excluded. For each time point the lateral displacement in y-direction was determined with the previous frame to generate the best overlap, which yields the y-directed migration velocity of a cell population.

### Migration within micro-fabricated polydimethylsiloxane (PDMS) based devices

Generation of PDMS-based devices and detailed experimental protocols can be found elsewhere^15,31^. Briefly, photomasks were designed using Coreldraw X18, printed on a photomask (5” square quartz, 1μm resolution, JD Photo data), followed by a spin coating step using SU-8 2005 (3000 RPM, 30sec, Microchem, USA) and a prebake of 2 min at 95°C. The wafer was then exposed to ultra-violet light (90 – 105 mJ/cm^2^ on an EVG mask aligner). After a post-exposure bake of 3 min 95°C, the wafer was developed in PGMEA. A one-hour silanization with Trichloro(1H,1H,2H,2H-perfluorooctyl)silane was applied to the wafer. The devices were made with a 1:10 mixture of Sylgard 184 (Dow Corning). Air bubbles were removed with a desiccator. The PDMS was cured overnight at 85°C. Micro-devices were attached to ethanol cleaned coverslips after plasma cleaning for 1h at 85°C. Before introduction of cells, devices were flushed and incubated with complete medium for at least 1h. To visualize the chemokine gradient, similar sized fluorescent dextran conjugated to fluorescinisothiocyanate (FITC) was added to the chemokine solution (200μg/ml, 10kDa, Sigma) due to exhibiting similar diffusion characteristics^32^.

### *In vitro* under-agarose migration assay

To obtain humid migration chambers a 17mm plastic ring was attached to a glass bottom dish using Paraplast X-tra (Sigma-Aldrich) to seal attachment site. For under-agarose migration assay, 4% Ultra Pure Agarose (Invitrogen) in nuclease-free water (Gibco) was mixed with phenol-free RPMI-1640 (Gibco) supplemented with 20% FCS, 1x Hanks buffered salt solution pH 7.3 in a 1:3 ratio. Ascorbic acid was added to final concentration of 50μM and a total volume of 500μl agarose-mix was cast into each migration chamber. After polymerization, a 2mm hole was punched into the agarose pad and 2.5μg/ml CCL19 (R&D Systems) was placed into the hole to generate a soluble chemokine gradient. Outer parts of the dish were filled with water followed by 30-minute equilibration at 37°C, 5% CO_2_. The cell suspension was injected under agarose opposite of the chemokine hole to confine DCs between coverslip and agarose. Prior to acquisition, dishes were incubated at least two hours at 37°C, 5% CO_2_ to allow recovery and persistent migration of cells. During acquisition, dishes were held under physiological conditions at 37°C and 5% CO_2_.

### Immunofluorescence

For fixation experiments a round shaped coverslip was placed in glass bottom dish before casting of agarose and injection of cells. Migrating cells were fixed by adding prewarmed 4% Para-Formaldehyde (PFA) diluted in cytoskeleton Buffer pH6.1 (10mM MES, 150mM NaCl, 5mM EGTA, 5mM Glucose, 5mM MgCl2) directly on top of the agarose. After fixation, agarose pad was carefully removed using a coverslip-tweezer followed by 20min incubation of the coverslip in 0.5% Triton X-100 in PBS and three subsequent washing steps á 10min with Tris-buffered saline (TBS) containing 0.1% Tween-20 (Sigma). Samples were blocked to prevent unspecific binding by incubating 60min in blocking solution (5% BSA, 0.1% Tween-20 in TBS). Immunostainings were carried out consecutively by 2h incubation with rat monoclonal anti-alpha-tubulin (AbD serotec), mouse anti-phospho-Myosin light chain 2 (S19) (Cell signaling), mouse anti-gamma tubulin (Sigma) or rabbit anti-acetylated alpha-tubulin (Sigma). Followed by 3×10min washing with blocking solution and 30min incubation using Alexa Fluor^®^ 488-AffiniPure F(ab’)_2_ or Alexa Fluor^®^ 647-AffiniPure F(ab’)_2_ Fragment IgG (H+L) (both Jackson Immuno) secondary antibodies. After incubation washing was done at least three times á 5min. Samples were conserved in non-hardening mounting medium with DAPI (VectorLaboratories) and stored at 4°C in the dark.

### Immunodetection of whole cell lysates

3×10^5^ cells were serum starved for 1h followed by drug treatment. After harvesting, cell pellet was snap frozen and lysed using RIPA buffer (Cell Signaling) to which 1mM Phenylmethylsulfonylfluoride was added prior to usage. Samples were supplemented with LDS Sample Buffer and Reducing agent (both Invitrogen) and incubated for 5min at 90°C before loading on pre-cast 4-12% Bis-Tris acrylamide gel (Invitrogen). Subsequently, samples were transferred to Nitrocellulose membrane using iBlot system (Invitrogen) and blocked for 1h in 5% bovine serum albumin in TBS containing 0.01% Tween-20. For whole cell lysate protein detection following antibodies were used: rabbit anti phospho-Myosin Light Chain 2 (S19) (1:500), rabbit anti Myosin Light Chain 2 (1:500), rabbit anti GEF-H1 (the mammalian homologue of Lfc) (1:500), rabbit anti phospho-ERM (1:500), rabbit anti ERM (1:500, all Cell Signaling), mouse anti glyceraldehyde 3-phosphate dehydrogenase (GAPDH) (1:1000, BioRad). As secondary antibodies Horseradish Peroxidase (HRP) Conjugated Anti-rabbit and anti-mouse IgG (H + L) antibodies were used in 1:5000 dilutions and enzymatic reaction was started by addition of chemoluminescent substrate for HRP (Super Signal West Femto). Chemoluminescence was acquired using a VersaDoc imaging system (BioRad). Western blot signals were quantified manually by normalization to input values and subsequent comparison of each treatment to signal intensity of steady-state level (i.e. control sample).

### Flow cytometry

Before staining, 1-2×10^6^ cells were incubated for 15 min at +4°C with blocking buffer (1xPBS, 1% BSA, 2mM EDTA) containing 5mg/ml α-CD16/CD32 (2.4G2, BD Biosciences). For surface staining, cells were incubated for 30 min at 37°C with conjugated monoclonal antibodies (mAbs; mouse α-CCR7-PE (4B12), rat α-mouse I-A/I-E-eFluor450 (M5/114.15.2), hamster α-mouse CD11c-APC (N418) diluted at the recommended concentration in blocking buffer. Flow cytometry analysis was performed on a FACS CANTO II flow cytometer (BD Biosciences).

### Pharmacological inhibitors

For perturbation of cytoskeletal and myosin dynamics we used final concentrations of 300nM Nocodazole and 10μM Y27632 (all purchased from Sigma Aldrich). Nocodazole was dissolved in dimethylsulfoxide (DMSO; Sigma Aldrich) and Y27632 in poly-buffered saline. Control samples were usually treated with 1:1000 DMSO if not indicated differentially.

### Fluorescent reporter constructs

Generation of a C-terminal eGFP fusion construct of Lfc was carried out by amplifying Lfc from DC cDNA using a NotI restriction site containing forward (5’ ATATGCGGCCGCAATCTCGGATCGAATCCCTCACTCGCG 3’) and reverse (5’ ATATGCGGCCGCTTAGCTCTCTGAAGCTGTGGGCTCC 3’) primer pair. After NotI digestion, Lfc was cloned into a pcDNA3.1 backbone containing eGFP (Express Link™ T4 DNA-Ligase). Correct sequence and orientation of clones was verified by sequencing (Eurofins). The fluorescent plasmid DNA reporter construct coding for EB3-GFP was a kind gift of V. Small (IMBA, Austria). M. Olson (Beatson Institute) generously provided MLC constructs (either fused to eGFP or RFP)^33^ and EMTB-3xmCherry constructs were a kind gift of (W. M. Bement, University of Wisconsin)^34^. Gateway cloning technology™ was employed to generate lentivirus from plasmid DNA constructs. Briefly, corresponding DNA segments were amplified using primers containing overhangs with *att*B1 and *att*B2 recombination sites on the 3’- and the 5’-end respectively. In order to obtain an EMTB fusion construct carrying a single mCherry tag, the PCR product was size separated via gel electrophoresis and only the fragment of corresponding size (EMTB: 816bp, mCherry: 705bp) was further processed. Gel purified PCR fragments were inserted into pcDNA221 entry vectors (Invitrogen) via BP recombination reaction, generating the entry clone. Expression clones were obtained by carrying out the LR recombination reaction between entry clone and pLenti6.3 destination vector (Invitrogen). Lentivirus production was carried out by co-transfecting LX-293 cells (Chemicon) with the expression clone of interest in conjunction with pdelta8.9 (packaging plasmid) and pCMV-VSV-G (envelope plasmid) (plasmids were a gift from Bob Weinberg)^35^. The supernatant of virus-producing cells was harvested 72h after transfection, snap frozen and stored at −80°C after sterile filtration.

### Transgene delivery

To induce expression of fluorescently labeled proteins DCs were transfected according to manufacturer guidelines using nucleofector kit for primary T cells (Amaxa, Lonza Group). Briefly, 5×10^6^ were resuspended in 100μl reconstituted nucleofector solution, transferred to an electroporation cuvette and a total amount of 4μg plasmid DNA was added. Cells were transfected by using a protocol specifically designed for electroporating immature mouse DCs (program X-001). After transfection, cells were cultured in 60mm cell culture dishes in complete media and taken for experiments 24h post-transfection. Due to low transfection efficiency of primary cells, transfected cells were FACS sorted prior to experiment using FACS Aria III (BD Biosciences).

### Luminometric RhoA activity assay

RhoA activities were determined using G-LISA™ RhoA Activation Assay Biochem Kit™ (Cytoskeleton) according to the manufacturer’s instructions. Briefly, 4×10^5^ mature BMDCs were lysed in 70μl RIPA buffer (Cell Signaling) and protein concentration determined using the Precision Red™ Advanced Protein Assay Reagent (Cytoskeleton). Respective samples were treated with 300nM Nocodazole for 15 min before lysis. All samples were adjusted to a final protein concentration of 0.5mg/ml. Luminescence signals were measured using a microplate photometer at 600nm. Wells containing lysis buffer only were used as reference blanks in all experiments.

### Microscopy

During all live cell imaging experiments cells were held under physiological conditions at 37°C, 5% CO2 in a humidified chamber. Low magnification bright field or DIC time-lapse acquisition was carried out using inverted routine microscopes (Leica), equipped with PAL cameras (Prosilica, Brunaby, BC) controlled by SVS-Visitek software (Seefeld, Germany) using 4x, 10x, 20x objectives or an inverted Nikon Eclipse widefield microscope using a C-Apochromat 20x/0.5 PH1 air equipped with a Lumencor light engine (wavelengths [nm]: 390, 475, 542/575). For high magnification live cell acquisition, either an Andor spinning disc confocal scanhead installed on an inverted Axio observer microscope (Zeiss), using a C-Apochromat 63x/1.2 W Korr UV-VIS-IR objective, or a total internal reflection (TIRF) setup consisting of an inverted Axio observer microscope (Zeiss), a TIRF 488/561 nm laser system (Visitron systems) and an Evolve™ EMCCD camera (Photometrics) triggered by VisiView software (Visitron) was chosen. Photo-activation experiments were conducted on an inverted Spinning disc microscope (iMic) using a 60x/1.35 Oil objective. TAMRA stained DCs were either untreated or treated with 10μM PST-1 in the dark and recorded using a 561nm laser line in 2-second intervals. Photoactivation was carried out on directionally migrating cells using a 405 nm laser line (pixel dwell time: 10 ms, interval: 40 sec). FRAP calibration was carried out on separate samples before each experiment. Acquisition of fixed samples (*in situ* ear crawl in and immunofluorescence samples) was carried out using an upright confocal microscope (LSM700, Zeiss) equipped with a Plan-Apochromat 20x/1.0 W DIC (UV) VIS-IR or a Plan-Apochromat 63x/1.4 Oil objective.

### Statistics

All boxes in Box-Whisker plots boxes extend from 25^th^ to 75^th^ percentile and whiskers span minimum to maximum values. Graphs represent pooled data of several cells (n) from independent biological experiments (N) as mentioned in the figure legends. Individual experiments were validated separately and only pooled if showing the same trend. For representation of frequencies, bar charts depict mean values from several independent biological experiments (N) ± S.D. Statistical analysis was conducted out using GraphPad Prism.

---

**Supplementary Figure 1.**
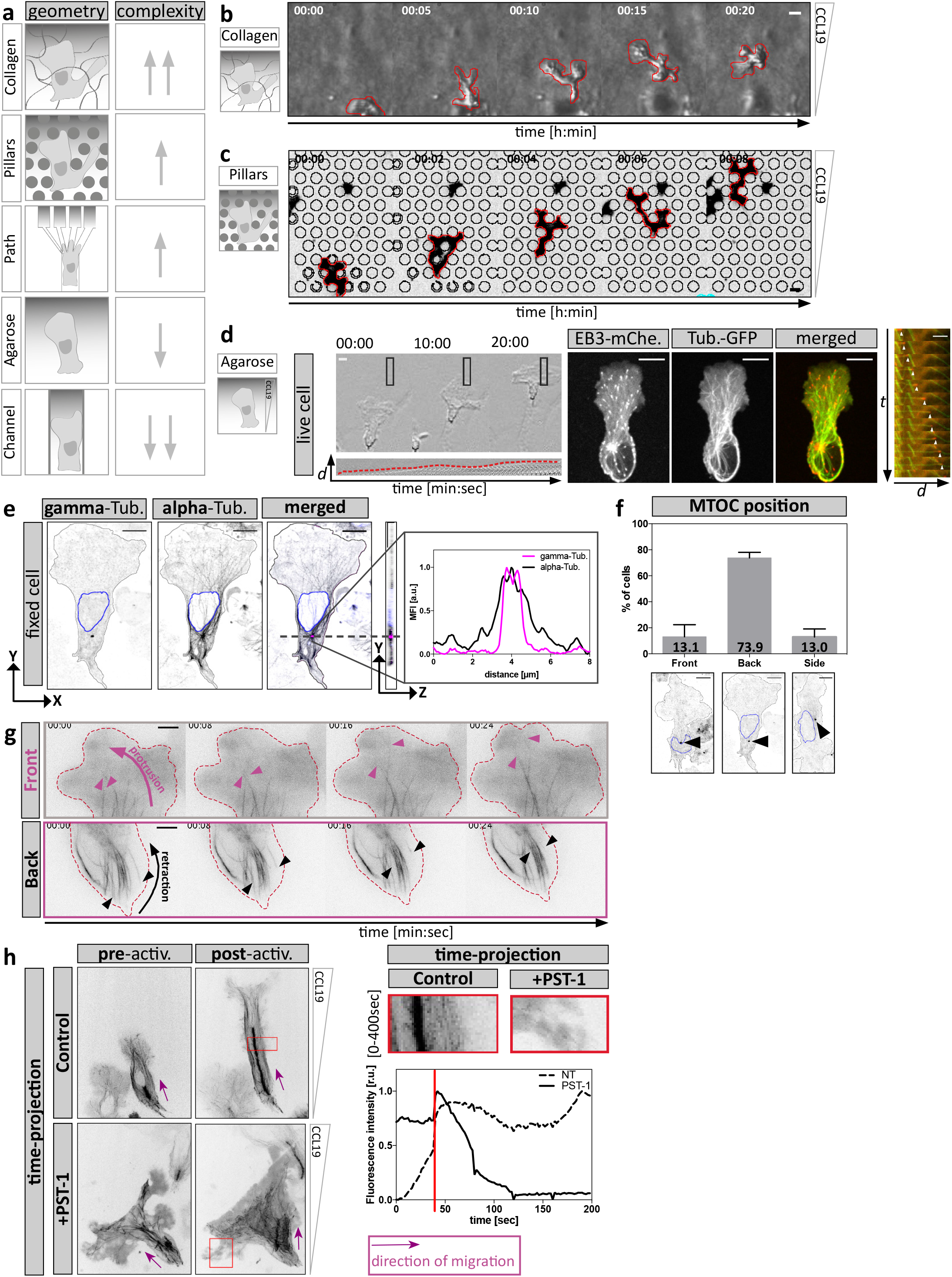
DC migration within diverse matrices to study the role of the MT cytoskeleton during cell migration. **a**, Schematic representation of migration assays used in this study. Assays range from highly complex (top) and relatively uncontrollable geometries to very simple and precisely controllable PDMS-based structures (bottom). Complexity of the geometrical confinement correlates with dynamic shape changes of cells. Upward-facing arrows indicate high geometrical complexity and cell shape changes respectively. Downward-facing arrows indicate low complexity. **b**, Cell shape changes of a DC migrating in a collagen matrix along a soluble CCL19 gradient. **c**, Dynamic cell shape changes are recapitulated during migration within a defined array of PDMS-based pillar structures. **d**, Left panel: Cells migrating under agarose display a protrusive lamellipodium (lower panel: montage of boxed area) followed by a contractile trailing edge. Scale bar, 10μm. Middle panel: EB3-mCherry localizes to the plus tips of tubulin-GFP decorated MT filaments. Shown is a double-reporter DC migrating under agarose along a soluble CCL19 gradient. Scale bar, 10μm. Right panel: EB3-mCherry faithfully tracks growing MT filaments during DC migration. White arrowhead highlight the localization of EB3 signal at the tip of polymerizing tubulin filaments as the cell advances. Scale bar 5μm. **e**, MT nucleation from centrosomal origin determined by alpha- and gamma-tubulin staining. Right panel shows line scan of signal intensities along purple line in merged image. Scale bar, 10μm. **f**, Determination of MTOC position by alpha- and gamma-tubulin staining with respect to the nucleus. Mean ± S.D. of n = 256 cells from N = 3 experiments. **g**, Time-course analysis of MT filament dynamics of migrating DCs expressing EMTB-mCherry. Upper panel indicates leading edge area. The purple arrow represents membrane protrusion and the purple arrowheads represent elongating MT filaments. Lower panel indicates trailing edge area in which black arrow represents membrane retraction and black arrowheads MT filament depolymerization. Red dashed line indicates cell edges. Scale bar, 10μm. **h**, EB3-mCherry localization of control or PST-1 treated cells migrating under agarose along a soluble CCL19 gradient. The red box indicates photo-activated area magnified on the right. Magnified regions show time projection of EB3-mCherry intensities after local photo-activation. Lower panel indicates fluorescence intensity evolution upon photo-activation of control or PST-1 treated cells. The red line highlights the time point of initial photo-activation. Purple arrows indicate the direction of cell migration.

**Supplementary Figure 2.**
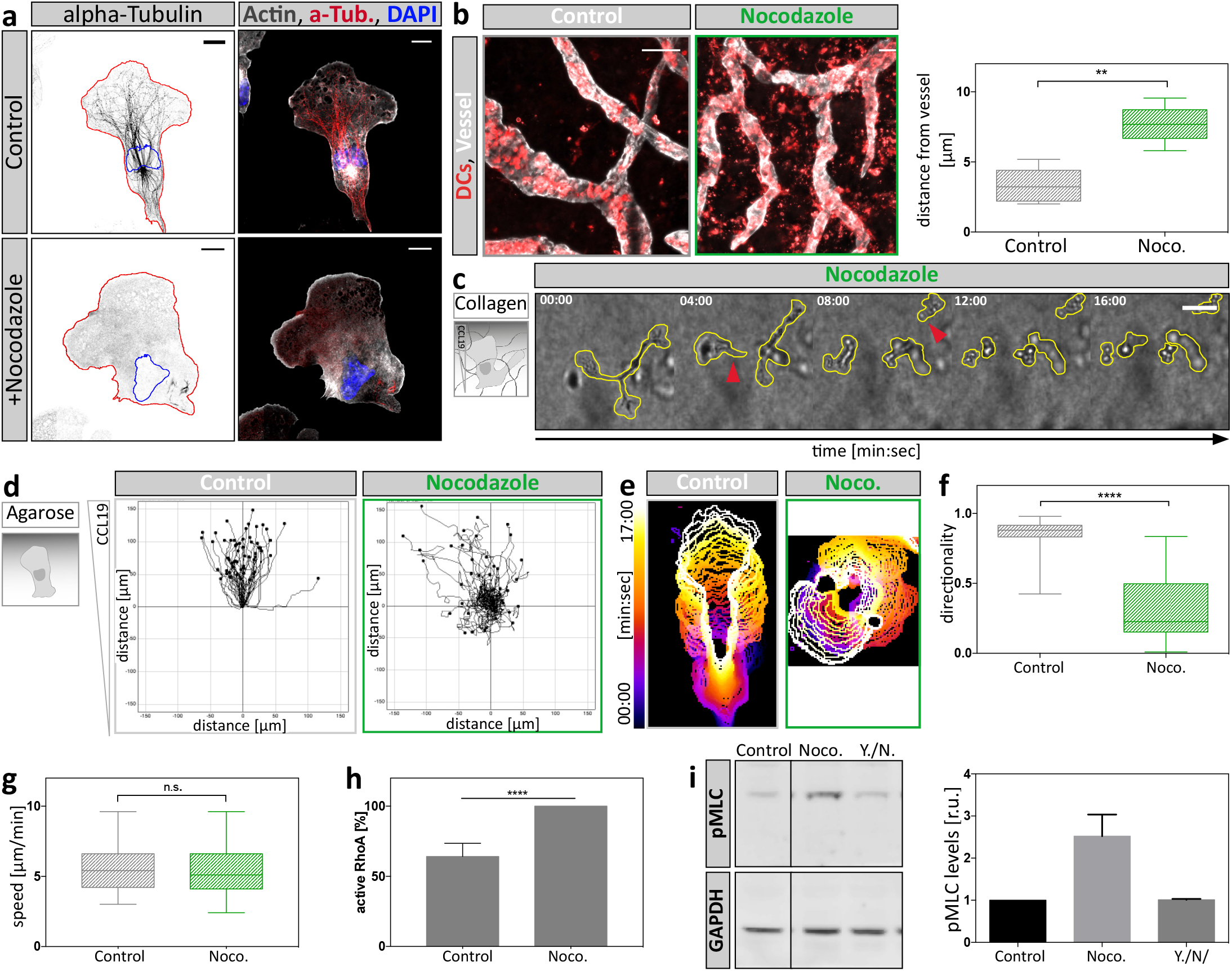
Perturbation of the MT cytoskeleton affects DCs migration on multiple levels. **a**, Non-treated control or nocodazole-treated cells migrating under agarose towards a CCL19 gradient were fixed and stained for endogenous distribution of alpha-tubulin and F-actin. Scale bar 10μm. **b**, *In situ* migration of endogenous DCs on a mouse ear sheet. Z-projections of separated ear sheets upon control conditions or nocodazole treatment. Lymphatic vessels were stained for Lyve-1, DCs for MHC-II. Mean distance from lymphatic vessels of endogenous DCs was determined 48h after ear separation (right panel). Per condition, four mouse ears with two fields of view were analyzed. Boxes extend from 25^th^ to 75^th^ percentile. Whiskers span minimum to maximum values. ** P ≤ 0.01. Scale bar, 100μm. **c**, Nocodazole-treated DC migrating in a collagen gel towards a soluble CCL19 gradient. Yellow line outlines cell shape. Red arrowheads indicate loss of cellular coherence. Scale bar, 100μm. **d**, Individual cell migration trajectories of cells migrating under agarose upon control conditions and nocodazole treatment of n = 58 cells (Control) and n = 52 cells (Noco.) from N = 4 experiments. **e**, Individual cell outlines over time upon control or nocodazole-treated conditions. Note the stable polarization of control cells contrasting the oscillatory protrusion dynamics of nocodazole-treated cells. **f**, Directionality during migration under agarose of n = 50 cells per condition from N = 4 experiments was assessed by comparing accumulated-with euclidean-distance of manually tracked cell trajectories. Boxes extend from 25^th^ to 75^th^ percentile. Whiskers span minimum to maximum values **** P ≤ 0.0001. **g**, Migration speed during migration under agarose along a soluble CCL19 gradient upon control conditions and nocodazole treatment of n = 50 cells per condition from N = 4 experiments. Boxes extend from 25^th^ to 75^th^ percentile. Whiskers span minimum to maximum values. **h**, Levels of active RhoA upon MT depolymerization with nocodazole determined by luminometry. RhoA activity levels were normalized to nocodazole-treated samples. Plotted is mean ± S.D. from N = 3 experiments. **** P ≤ 0.0001. **i**, Levels of MLC phosphorylation determined by Western Blot analysis. Cells were treated with the indicated compounds (DMSO, nocodazole or Y27632 together with nocodazole (Y./N.)). Mean fluorescence intensity of pMLC was normalized to GAPDH signal and shown as fold increase relative to DMSO control ± S.D in right panel. Blots are representative of N = 3 experiments.

**Supplementary Figure 3.**
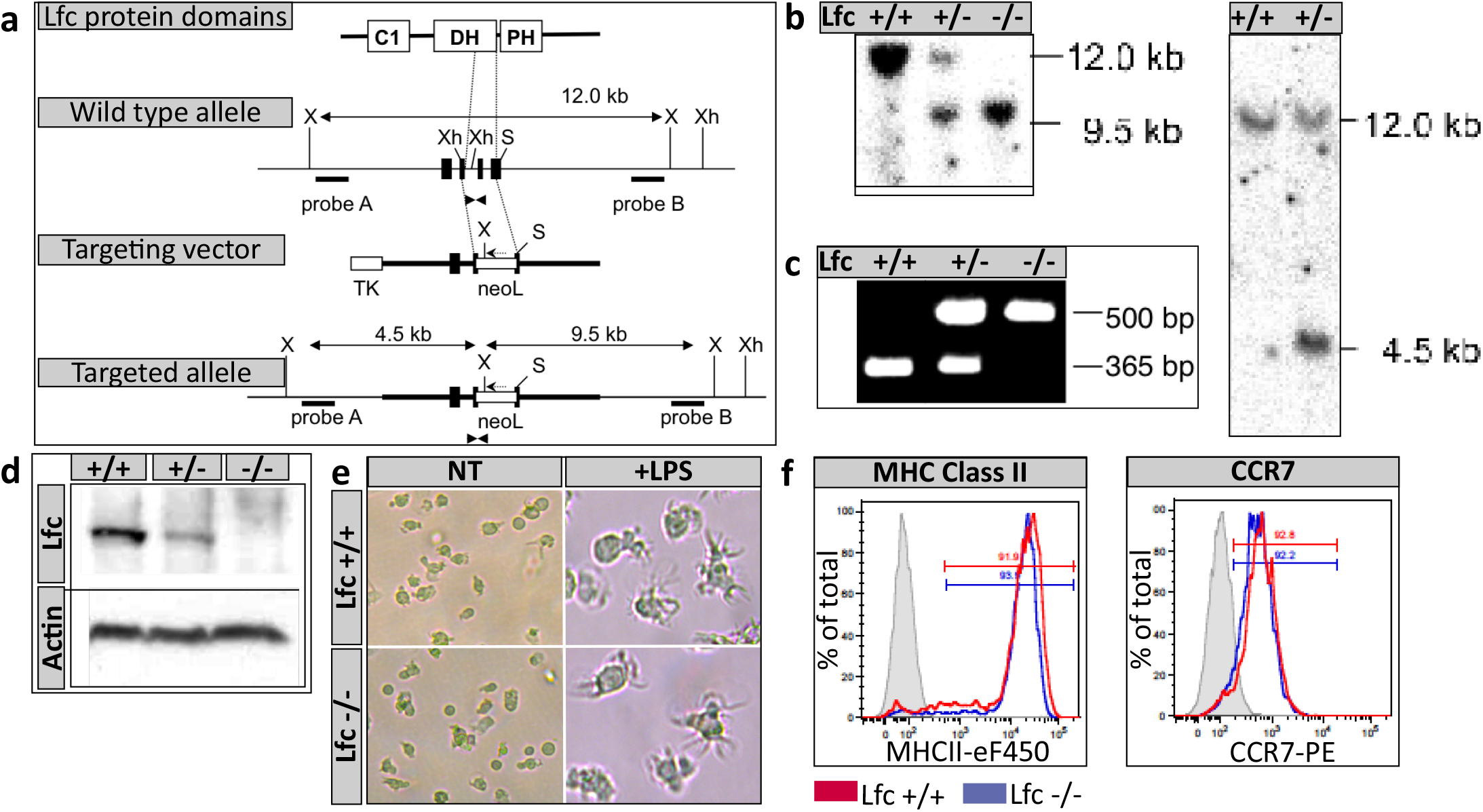
Generation of a Lfc^-/-^ mouse line. **a**, Integration of the Lfc targeting vector into genomic locus. Black boxes represent exons. The neo-lox P cassette was cloned in reverse orientation into two, replacing a *SmaI-XhoI* segment. Locations of primers used for PCR are indicated with triangles. Probes A and B were used for Southern blot detection of short and long arms, respectively. S, *Sma*I; Xh, *Xho*I; X*, Xba*I: N, *Nhe*I. **b**, Southern blot analysis. Genomic DNA from Lfc^*+/+*^, Lfc^*+/-*^ and Lfc^-/-^ mice was digested with *Xba*I and hybridized with probes B (left panel) and genomic DNA from Lfc^*+/+*^ and Lfc^*+/-*^ embryonic stem cells were hybridized with probe A (right panel). **c**, PCR analysis of tail DNA from Lfc^*+/+*^, Lfc^+/-^ and Lfc^-/-^ mice. Locations of primers used for PCR are indicated with triangles in **a**. **d**, Immunoblot analysis of total thymus cell lysates probed for Lfc protein content. **e**, Cell morphologies of immature (NT) and mature (+LPS) Lfc wildtype (upper-lane) and Lfc-deficient (lower-lane) littermate DCs. Note the presence of multiple veils in both LPS-treated samples. **f**, DC differentiation markers (MHC-II and CCR7) of Lfc^*+/+*^ (blue line) and Lfc^-/-^ (red line) littermate DCs compared to unstained cells (grey peak).

**Supplementary Figure 4.**
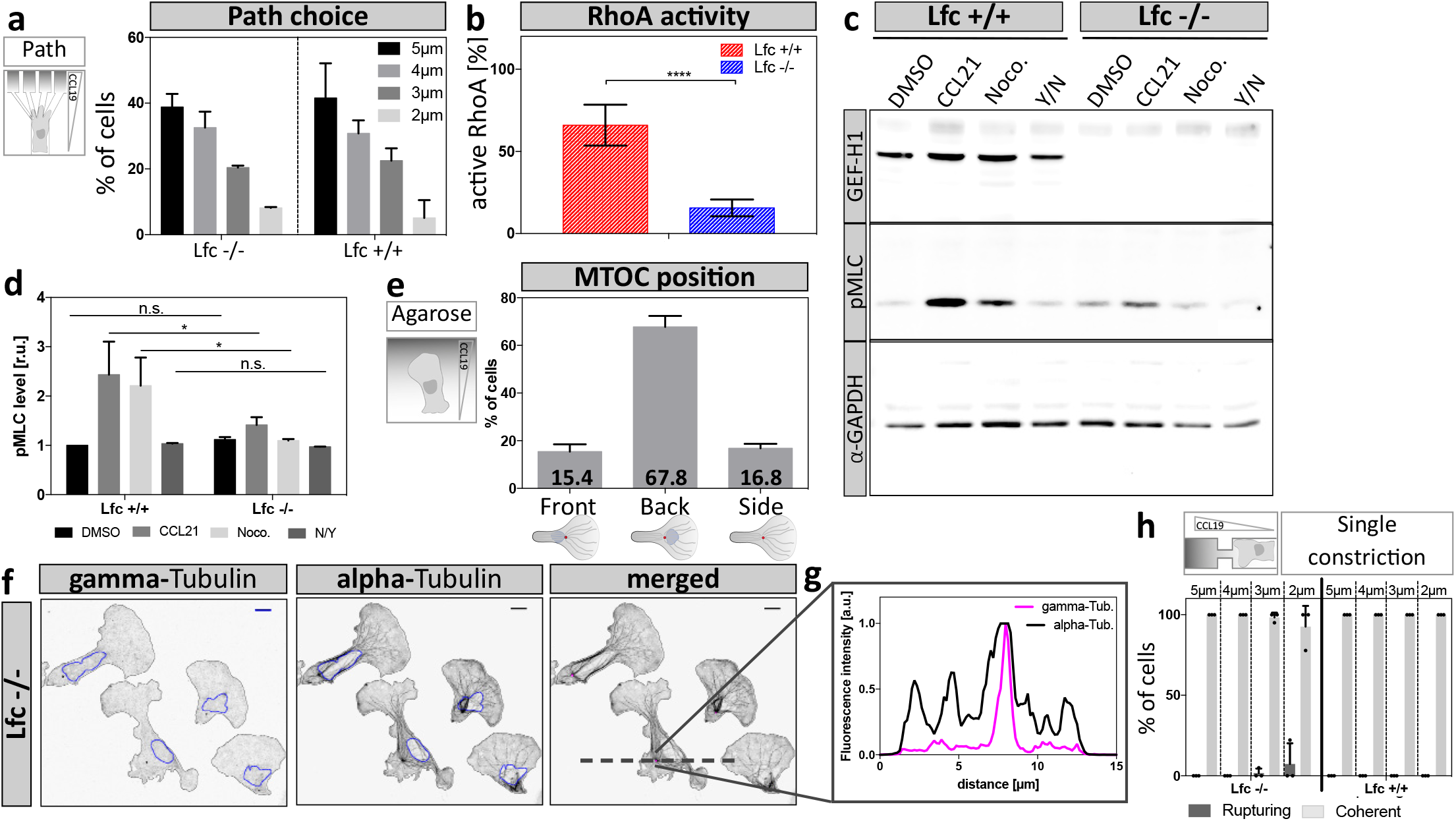
Lfc^*-/-*^ DCs exhibit reduced contractile responses. **a**, Path choice preference of Lfc^*+/+*^ and Lfc^-/-^ DCs migrating within a path choice assay. Shown are mean frequencies of Lfc^-/-^ (n = 49 cells of N = 2 experiments) and Lfc^*+/+*^ (n = 79 cells of N = 3 experiments) DCs. **b**, Levels of active RhoA of Lfc^*+/+*^ and Lfc^-/-^ cells was determined by luminometry showing mean intensities ± S.D. from N = 3 experiments. **** P ≤ 0.0001. **c**, Levels of MLC phosphorylation in Lfc^*+/+*^ and Lfc^-/-^ DCs assessed by Western Blot analysis. Cells were treated with the indicated compounds (DMSO, CCL21, nocodazole, Y27632 together with nocodazole). **d**, Mean fluorescence intensity of phospho-MLC was normalized to GAPDH signal and shown as fold increase relative to DMSO control ± S.D. Blots are representative of N = 3 experiments. **e**, Centrosome localization in Lfc^-/-^ DCs migrating under agarose assessed by alpha- and gamma-tubulin co-staining (n = 117 cells from N = 2 experiments). **f**, MT nucleation from centrosomal origin as determined by alpha- and gamma-tubulin co-staining. Scale bars, 10μm. **g**, Intensity line scans across the highest gamma-tubulin signal along the left-right axis (dashed line in **f**). The purple line indicates gamma-tubulin signal intensity. The black line indicates alpha-tubulin signal distribution. **h**, Frequency of cell rupturing events of Lfc^*+/+*^ (n = 73 cells, N = 3 experiments) and Lfc^-/-^ (n = 128 cells, N = 3 experiments) DCs while migrating within single constriction containing microchannels.

**Supplementary Figure 5.**
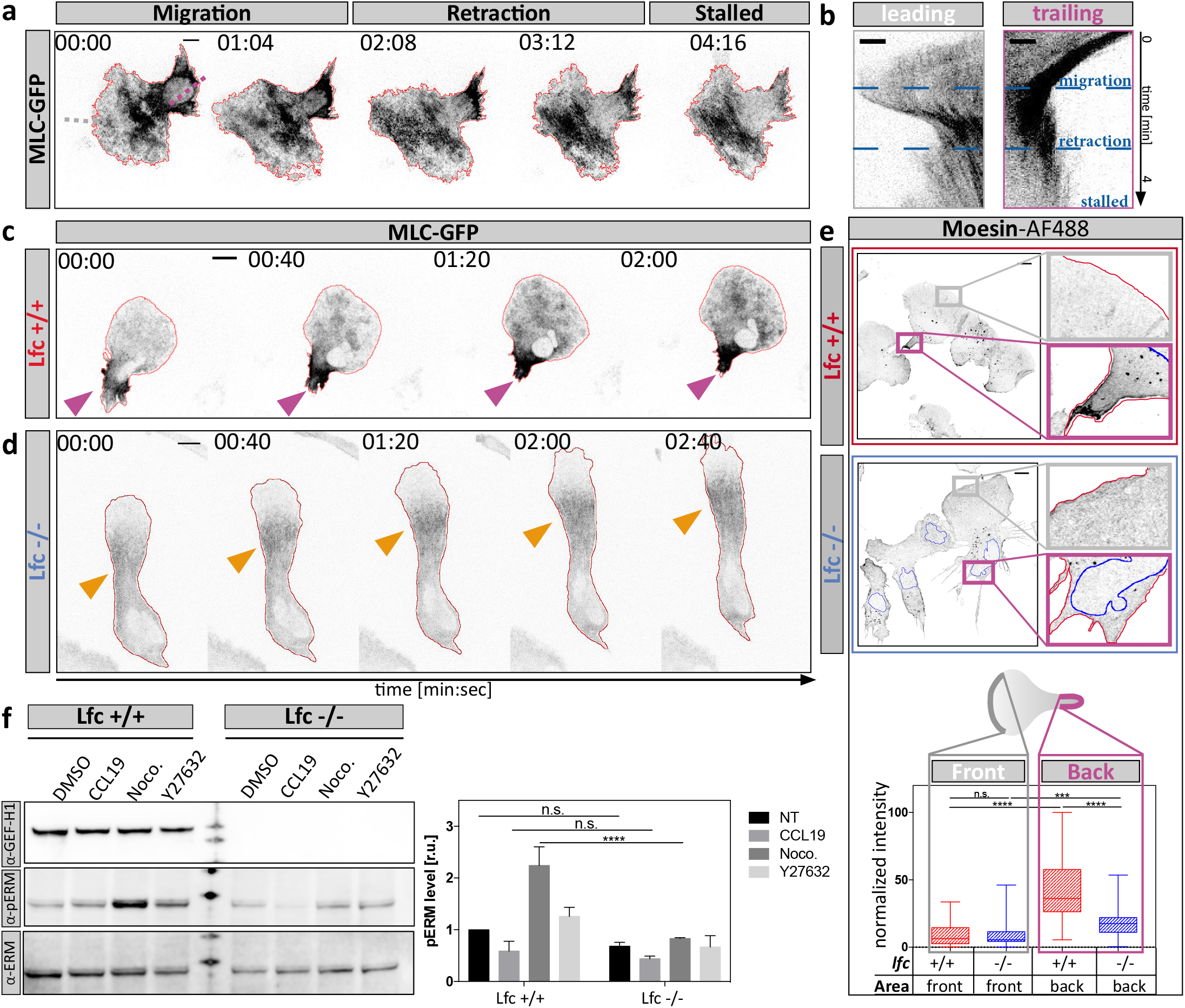
Aberrant spatiotemporal MLC accumulation and moesin localization in Lfc^-/-^ DCs. **a**, Time-lapse montage of a MLC-GFP expressing DC migrating under agarose towards a soluble CCL19 gradient. A cycle of migration, retraction, and pausing is shown. Scale bar, 10μm. Dotted lines indicate positions further analyzed by Kymograph in **b**. **b**, Leading edge kymograph was derived from grey dotted line in leading edge region of **a**. Trailing edge kymograph was derived from purple dotted line in trailing edge region of **a**. Note the absence of MLC accumulation in leading edge areas and the presence of trailing edge MLC accumulation during migration Scale bar, 5μm. **c**, Time-lapse sequence showing spatiotemporal MLC accumulation of a Lfc^*+/+*^ DC and **d**, a Lfc^-/-^ DC. Purple arrowheads highlight trailing edge MLC accumulation, orange arrowheads indicate central MLC accumulation. Scale bars, 10μm. **e**, Quantitative morphometry of Moesin in fixed migratory Lfc^*+/+*^ (red) and Lfc^-/-^ (blue) DCs. Lower panel: Quantification of fluorescence intensity in leading versus trailing edge regions of Lfc^*+/+*^ (red) and Lfc^-/-^ (blue) DCs of n = 55 cells per condition from N = 3 experiments. Boxes extend from 25^th^ to 75^th^ percentile. Whiskers span minimum to maximum values. *** P ≤ 0.001, **** P ≤ 0.0001. Scale bars, 10μm. **f**, Protein levels of phospho-ERM in Lfc^*+/+*^ and Lfc^-/-^ DCs assessed by Western Blot analysis. Right panel: Quantification of pERM levels upon treatment with DMSO, CCL19, nocodazole or Y27632. Mean fluorescence intensity of pERM signal was normalized to total ERM signal and shown as fold increase relative to Lfc^*+/+*^ DMSO control ± S.D. of N = 3 experiments.

**Supplementary Figure 6.**
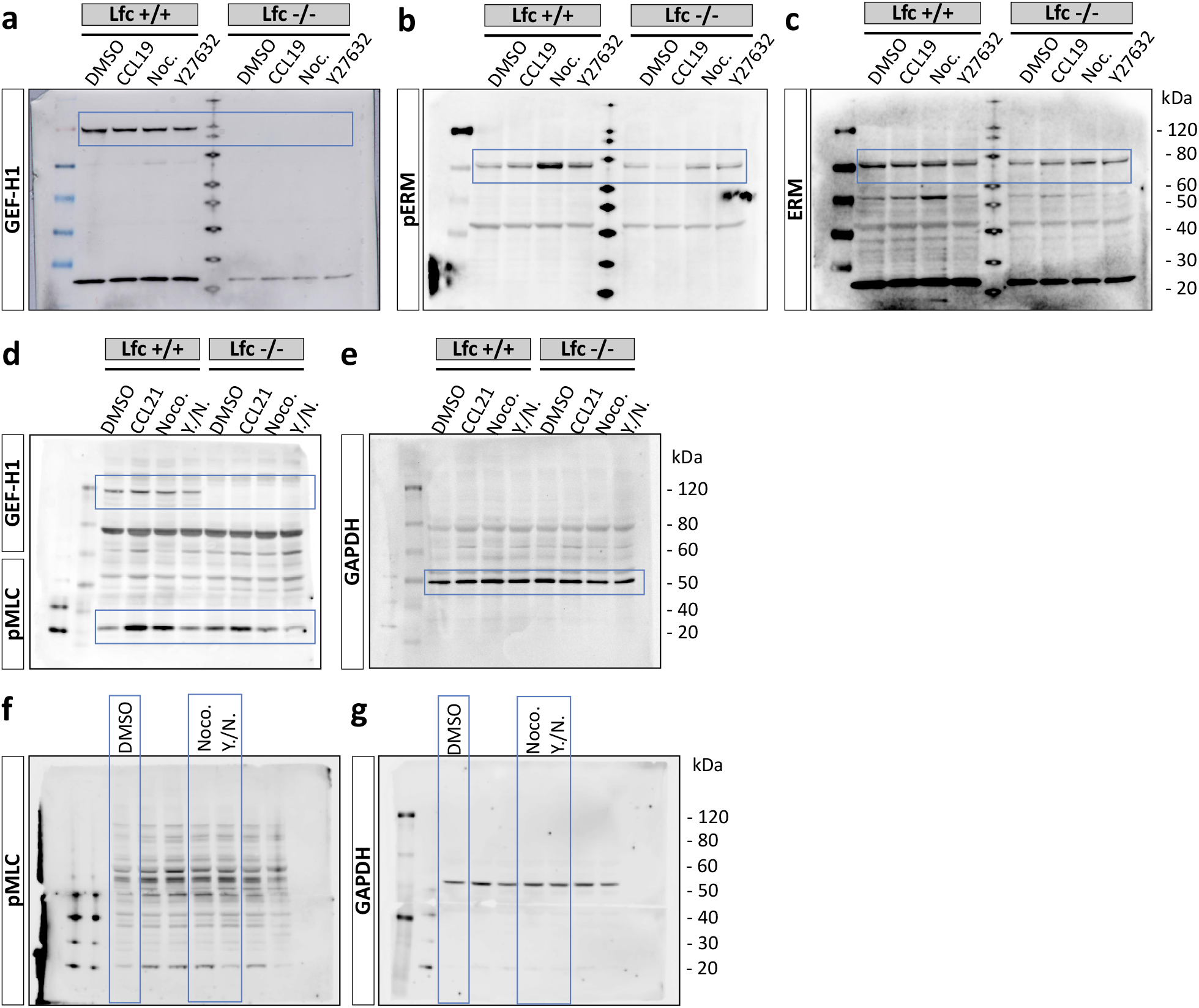
Unprocessed western blot scans. **a**, Raw image of blot probed against GEF-H1. Blue boxed region indicates area shown in Supplementary Fig. 5f. **b**, Raw image of blot probed against phospho-ERM proteins. Blue boxed region indicates area shown in Supplementary Fig. 5f. **c**, Raw image of blot probed against total ERM protein. Blue boxed region indicates area shown in Supplementary Fig. 5f. **d**, Raw image of blot probed against GEF-H1 and phosho-MLC. Blue boxed regions indicate areas shown in Supplementary Fig. 4c. **e**, Raw image of blot probed against GAPDH. Blue boxed region indicates area shown in Fig. Supplementary 4c. **f**, Raw image of blot probed against phosho-MLC. Blue boxed regions indicate areas shown in Supplementary Fig. 2i. **g**, Raw image of blot probed against GAPDH. Blue boxed region indicates area shown in Supplementary Fig. 2i.

## Supplementary Movie Legends

**Supplementary Movie 1. MT dynamics during path finding within a pillar array.** An EB3-mCherry expressing reporter cell was acquired while migrating within a complex 3D pillar array towards a soluble CCL19 gradient in 2sec intervals on an inverted spinning disc microscope. Scale bar, 10μm.

**Supplementary Movie 2. Polarized MT dynamics in migratory DCs.** DC is expressing EMTB-mCherry. Migration during 2D confinement under agarose was acquired in 2sec intervals using a TIRF setup. For representation, the signal was inverted after the acquisition. The upper panel shows the protruding leading edge, in which grey arrowheads indicate elongating MT filaments. The lower panel shows retracting trailing edge of the same cell in which purple arrowheads highlight MT shrinking events. Time in [min:sec]. Scale bar, 5μm.

**Supplementary Movie 3. Induced MT depolymerization locally activates the contractile module.** TAMRA stained DCs migrating under agarose were recorded every 2sec on an inverted spinning disc microscope and locally photo-activated (red box) every 40sec using a 405nm laser line. Cells were either untreated (upper panel) or treated with PST-1 (lower panel). Time in [min:sec]. Scale bar 10μm.

**Supplementary Movie 4. MTs promote cell coherence during migration in complex environments.** DCs were either non-treated (control) or treated with the indicated compounds (nocodazole or double treatment using Y27632 together with nocodazole) and recorded while migrating within a path choice assay towards a soluble CCL19 gradient in 60sec intervals. Note that under all conditions cells insert multiple protrusions into different channels when reaching the junction point (black arrowheads). Red arrowheads highlight rupturing events and loss of cellular coherence only observed in drug-treated cells. Time in [min:sec]. Scale bar, 10μm.

**Supplementary Movie 5. Perturbation of MT and myosin dynamics impairs DC migration in complex scaffolds.** Mature DCs migrating along a soluble CCL19 gradient within a 3D collagen matrix. Shown are separately acquired bright field movies of control- (DMSO), nocodazole-treated and double-treated cells using Y27632 and nocodazole (Y/N) reconstructed in a single file. Images were acquired every 60sec for 5h and are represented as single movie in 4min intervals. Time in [min:sec]. Scale bar, 100μm for representative movie of bulk cell movement, scale bar, 10μm for movie showing single cell dynamics.

**Supplementary Movie 6. Perturbation of MT and myosin dynamics permits DC migration within simple linear microenvironments.** Mature DCs migrating along a soluble CCL19 gradient within a straight microchannel. Shown are separately acquired bright field movies of non-treated, nocodazole-treated and double-treated cells using Y27632 and nocodazole cells reconstructed in a single file. Images were acquired in 20sec intervals for 5h. Note the frequent directional oscillations of nocodazole only treated cells. Time in [min:sec]. Scale bar, 10μm.

**Supplementary Movie 7. Microtubules mediate retraction of supernumerary protrusions via Lfc.** Lfc^*+/+*^ and Lfc^-/-^ DCs were recorded while migrating within a path choice assay towards a soluble CCL19 gradient in 30sec intervals. Note that both genotypes insert multiple protrusions into different channels when reaching the junction point (black arrowheads). Red arrowheads highlight rupturing events and loss of cellular coherence only observed in Lfc-deficient cells. Time in [min:sec]. Scale bar, 10μm.

**Supplementary Movie 8. Lfc specifies myosin localization at the trailing edge.** Combined movies of MLC-GFP expressing Lfc^*+/+*^ DCs (left panel) and Lfc^-/-^ DCs (right panel) migrating under agarose along a soluble CCL19 gradient, acquired in 2sec intervals on an inverted spinning disc microscope. Magenta arrowhead indicates trailing edge MLC accumulation, which is absent in Lfc^-/-^ cells. Orange arrowhead highlights central MLC accumulation. Fluorescence signal was inverted for better visualization. Time in [min:sec]. Scale bar 10μm.

**Supplementary Movie 9. Lfc promotes DC migration within complex environments.** Mature DCs generated from Lfc^+/+^ and Lfc^-/-^ mice were embedded in a 3D collagen matrix. Migration along a soluble CCL19 gradient was acquired for 5h in 1min intervals. Images for each condition were subsequently reconstructed as a single file in 4min intervals. Time in [min:sec].

**Supplementary movie 10. Lfc regulates MT-mediated adhesion resolution.** Nocodazole-treated Lfc^*+/+*^ and Lfc^-/-^ DCs were acquired while migrating under agarose towards a soluble CCL19 gradient in 20 second intervals on an inverted cell culture microscope. Left panels show nocodazole-treated cells during adhesive migration. Note the loss of directionality in Lfc^*+/+*^ DCs and the pronounced elongation of Lfc^*-/-*^ DCs. Right panels show nocodazole effects during adhesion-independent migration on PEG coated coverslips. Note the persistent loss of directionality in Lfc^*+/+*^ DCs but the restored cell lengths of of Lfc^-/-^ DCs. Time in [min:sec]. Scale bar 100μm.

